# Specialized NADPH diaphorase membrane-related localizations in the brainstem of the pigeons (Columba livia)

**DOI:** 10.1101/663310

**Authors:** Yunge Jia, Wei Huo, Yinhua Li, Tianyi Zhang, Xinghang Wang, Xiaoxin Wen, Ximeng Xu, Haoran Sun, Xianhui Wu, Chenxu Rao, Zichun Wei, Zhenhua Zhai, Huibing Tan

## Abstract

NADPH diaphorase (N-d) is used to a histochemical identification of subgroup of neuronal cells. Beside regular intracellular N-d positivity, membrane-related positivity revealed as a specialized staining pattern in the pigeon brain stem. In the investigation of the nervous system of homing pigeons (Columba livia) with N-d staining, we found a specialized structure, which temporally was termed as N-d tubular glomerular body/structure or as T-J body related to the last name of authors. This N-d positive specialization constituted by tubular components bilaterally located in the medial to the lemniscus spinalis in the medulla oblongata. The tubular components were moderate staining. T-J body was a longitudinal oriented structure of 2400 μm with N-d staining. N-d positive tubular components were twisted and intermingled together. Beside the young adult pigeons, T-J body s were also consistently detected in the aged pigeons. Membrane-related staining were also detected in the other rostral nuclei in the brain stem. With discussion and review of related scientific literatures, T-J body was considered as a new anatomical structure or a new feature of the existent nucleus. In summary, beside N-d intracellular distribution, there were other three N-d membrane-related localizations: mini-aggregation, patch-aggregation, and arrangement along tubular unit.

## Introduction

NADPH diaphorase (N-d) is used to a histochemical identification of neuronal cells[1, 2] and also used to detect the neurodegenerative and neuropathological neurites[3–7]. Our research interesting is focused on the aging alterations in the lumbosacral spinal cord and dorsal column nuclei. Aging-related neurodegeneration are also detected in the pigeons[8]. The N-d positivity vary with neurons [9] and neuroglia[10] as well as blood vessel endothelium[11, 12]. Morphological feature of neuronal fibers or neurites are definitely distinguishable from neuroglia and blood vessels[11, 13]. The similar observation of the brain and the spinal cord of pigeons are well demonstrated in the anatomical morphology of N-d positivity [8, 14–16]. The gastrointestinal tract of Coturnix coturnix, an avian species is innervated by N-d-ergic neurons, as it is in mammals. Pigeon cloaca, a unique structure for avian is also innervated by N-d-ergic neurons [16]. In general, the N-d positive neurons in the pigeons are similar to the other species[8]. For neuronal cells, N-d histochemistry produces subgroup of positivity based on size of the somata, intensity of enzyme reaction (strong or light intensity), visible processes and less or un-visible process. Many neurons are visible for Golgi-like staining. Some of endothelia of blood vessels are stained with N-d histochemistry. Impression of the N-d staining is superiority to reveal specialized formation, such as aging-related N-d body and megaloneurite in the aged animals[4, 6–8]. However, recently, we found an uncommon N-d positive specialized structures featured with tubul-like components twisted and intermingled formation in the young adult pigeons. The aim of the investigation is to described the newly discovery of the anatomical structure.

## Methods and Materials

### Animals and Tissue Preparation

Experiments were performed using young adult pigeons (Columba Livia, one-year-old, n = 12) and aged pigeon (10~15-year-old, n=6). For age confirmation, every pigeon had identified number of leg ring. The pigeon owners had approved certification for breeding the pigeons and taken part in local pigeon club. Pigeons were housed in home-made facilities. Food and water were provided properly. All experimental procedures were approved by the Ethics Committee in Animal and Human Experimentation of the Jinzhou Medical University. All animals were anesthetized with pentobarbital sodium (45 mg/kg, i. p.). The chest cavity was opened and a cannula was inserted into the left ventricle. Perfusion was performed using 100–150 ml of 0.9% NaCl followed by 4% paraformaldehyde in 0.1M sodium phosphate buffer (PB, pH 7.4). The spinal cord and brain were rapidly removed and postfixed with 4% paraformaldehyde in 0.1 M phosphate buffer, left at 4□ for 2h and then placed in 30% sucrose for 48 h.

### Histochemical processing

Staining was performed using free-floating sections. The brains including medulla oblongata (n=8) were cut coronally into 40µm thick sections using a Leica CM1950 cryostat. The other four tissues were cut horizontally. Part of the slice was stained and examined by N-d histochemistry according to the previous study. Briefly, for N-d histochemistry, free-floating sections were incubated in PB containing Triton X-100 (0.3%), NADPH (1 mg/ml, Sigma) and nitroblue tetrazolium (1 mg/ml, Sigma) at 3□for 90 to 120 min. The reaction was stopped by the phosphate buffered saline (PBS, 0.01M). In order to distinguish membrane-related N-d locations to intracecullar locations, part of sections was stained in extra volume incubating solution with extended time for testing that the membrane-related N-d positivity was fixed the membrane location and not cause strong spread intracellular staining. The sections were examined and photographed under an optical microscope (Olympus BX35, camera DP80, Japan). Part of sections were counterstained with neutral red (0.5%). For Nissl staining, the frozen sections were stained with 0.1% cresyl violet for 10-20 min, rinsed with PBS, dehydrated in a graded alcohol series, cleared with xylene and mounted with neutral gum. The sections were observed using light microscopy.

### Measurement and statistics

Images were captured with a DP80 camera in an Olympus BX51 microscope (Olympus Shenyang, China). All Sections were observed under the light microscope. For specialized N-d positivity, one third sections selected from the brainstem in each animal were quantitated using Olympus image analysis software (Cellsens Standard, Olympus). A camera-lucida assistant function was used to draw the contour of observation.

## Results

In the brain stem, patterns of N-d neuronal positivity were sub-grouped based on size of the somata, intensity of enzyme reaction, processes visible and or un-stained processes (supplemental figure 1). We named the class of N-d staining as regular intracellular staining. Contract to these regular staining, especially Golgi-like staining, some N-d neuronal positivity revealed membrane-related location in the pigeon brain stem.

In the coronal sections, a moderate reactivity for N-d tubular glomerular structures symmetrically occurred in the medial to the lemniscus spinalis. We named the N-d tubular glomerular structures as T-J body or T-J complex which was named after the last name of corresponding author and first author (Figure 1). The word “body” referred to olivary body / olivary complex because the location of T-J body may be overlapped with superior olivary body or plexus Horsley. The term “glomerular” is referred to the glomerulus in olfactory bulb. The substructure components of the body were N-d positive tubular units. This N-d positive specialization constituted by tubular components bilaterally located in the medial to the lemniscus spinalis in the medulla oblongata. The diameter of the N-d tubular units was 11.46±0.58μm. In the coronal sections, these substructure units were twisted or intermingled together to form a glomerulus-like structure in the coronal transverse sections. The N-d tubular units distal to or toward the T-J body become solid fibers instead of tubular units (Figure 1, A2). The maximum diameter of the transection of glomeruli was arranged about 300-400μm. The glomeruli were reconstructed as a longitudinal structure by serial coronal sections. The length of T-J body was rostral-caudally oriented about 800-2400μm calculated through serial transections from rostral to caudal brainstem. The perikaryons could link to tubular components. Clear ramification was detected in the tubular components. However, regular dendritic tree hierarchy was not detected. There were N-d neurons scattered in the T-J body (Figure 2 B1). T-J body was also found in the aged pigeons (data not showed here). Nissl staining failed to reveal the N-d tubule glomerular body (Figure 2, F and F1). The sections counterstained with neutral read showed that membrane-related positivity occurred with the N-d formazans. In order to demonstrate the membrane-related N-d positive in the present investigation, we temporally termed the structure as peri-N-d-cytes, because we did not confirm the positive staining as regular N-d positive neurons. For distinguish membrane-related N-d locations to intracecullar locations, extended longer time staining did not cause the higher intensive intracellular staining (Figure 3). Next, in order to show perikaryons and membrane-related N-d locations, the counterstain sections with neutral red showed that the membrane-located formazans surrounded unreactive N-d perikaryons counterstained with intracellular neutral red (Figure4). Beside the N-d glomerulus, another pattern of peri-N-d-cytes could be detected in the rostral of the T-J body. These peri-N-d-cytes individually occurred with mini-aggregated N-d positivity formed peri-N-d-cytes (Figure 5). In further rostral, there were the other patched N-d positivity which occurred membrane-related locations and surround unstained perikaryons (Figure 6). There were also peri-N-d-cyte showed in the nucleus magnocellularis cochlearis (data not showed here).

**Figure 1.**
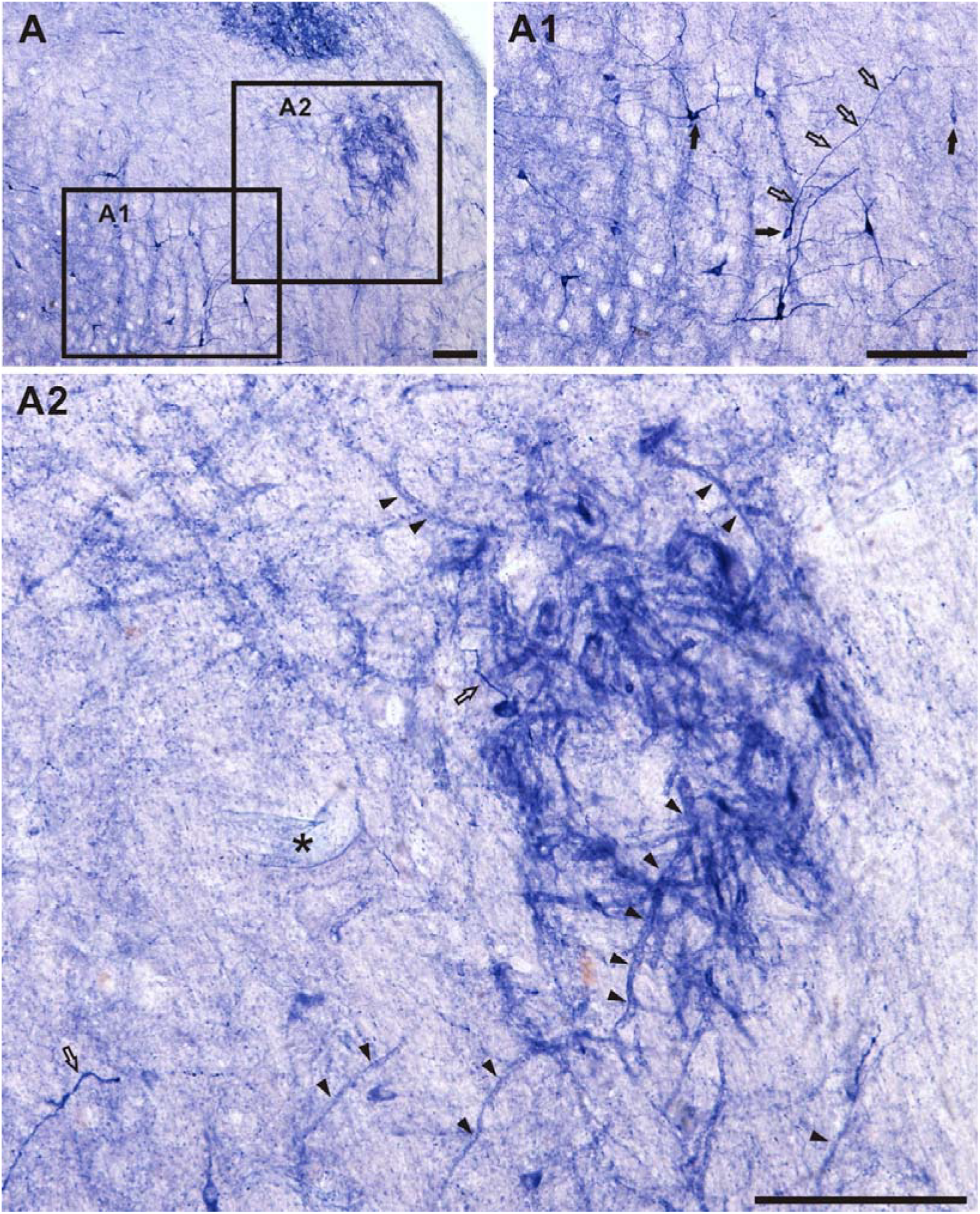
The N-d tubular glomerular structures of the coronal section: T-J body in the brainstem. A: N-d staining of the transection through the brainstem of the pigeon. A1: Neurons and fibers showed with higher magnification image from A. Arrow indicates neuron cell body. Open arrow indicates neuronal fibers. A2: Higher magnification image from A. The N-d tubular glomerular structure named as T-J body. Arrowhead indicate N-d tubular unit. Open arrow indicates neuronal fiber. Asterisk indicates N-d positive endothelium of blood vessel. Bar = 20μm

**Figure 2.**
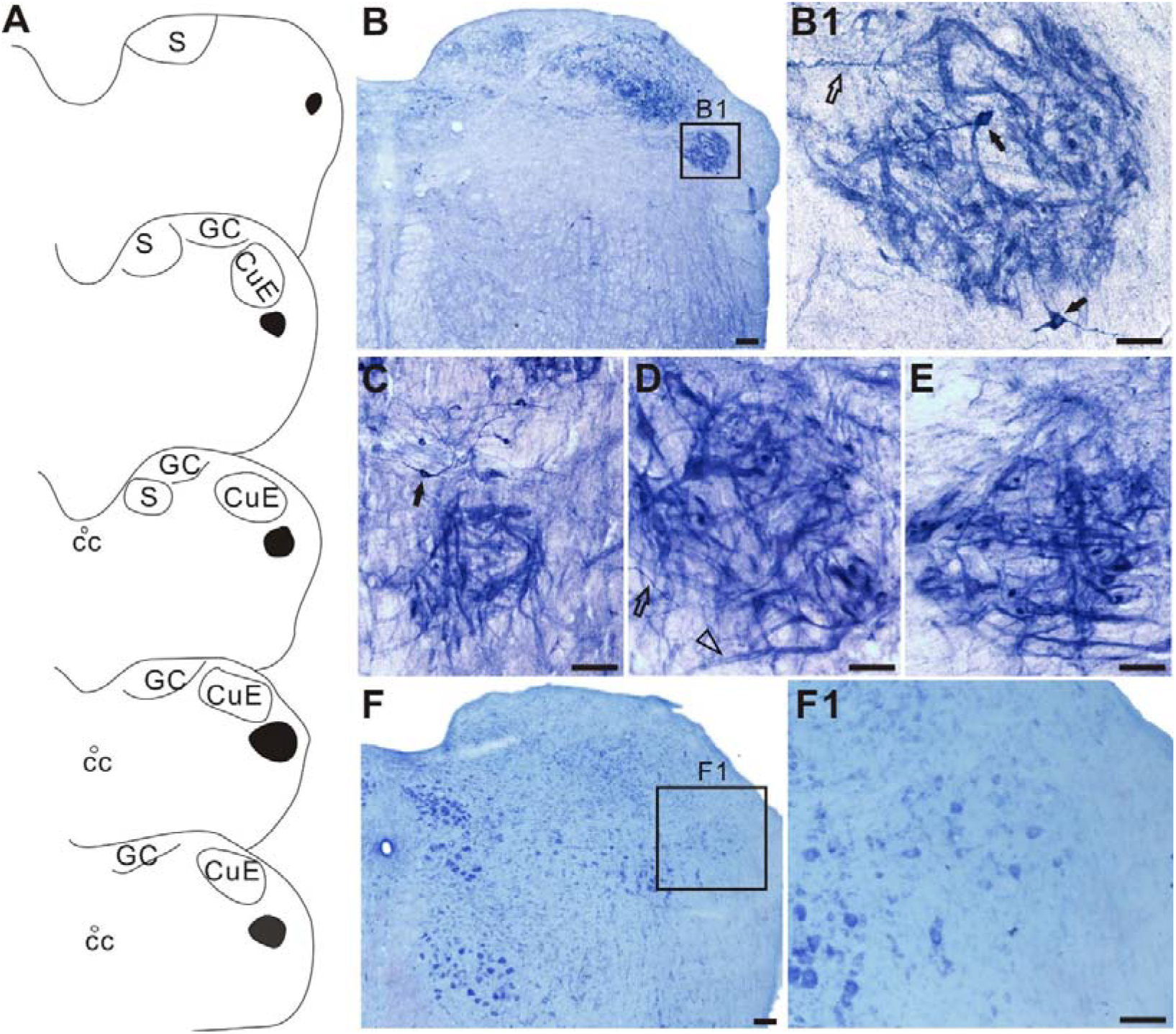
Location of T-J body and examples of T-J body in the brainstem. A: Drawings of structures of localization of T-J body. Black area indicates T-J body. cc: central canal; S: solitary nucleus; GC: nucleus gracilis et cuneatus; CuE: nucleus cuneatus externus. B: Low power of photo image of N-d positivity section. Box B1 indicates T-J body. B1: Higher magnification of B for T-J body. Arrow indicates built-in N-d neurons. C-E: The other three examples of T-J body. Open arrowheads indicate transverse of tubule-like units. Open arrow indicate regular N-d fibers. F: Nissl staining of the coronal transverse section in the medulla oblongata. F1 indicates the location of T-J body. F1: higher magnification of box F1 in F. Bar for B and F = 100μm. Bar for B1-E and F1 = 20μm

**Figure 3.**
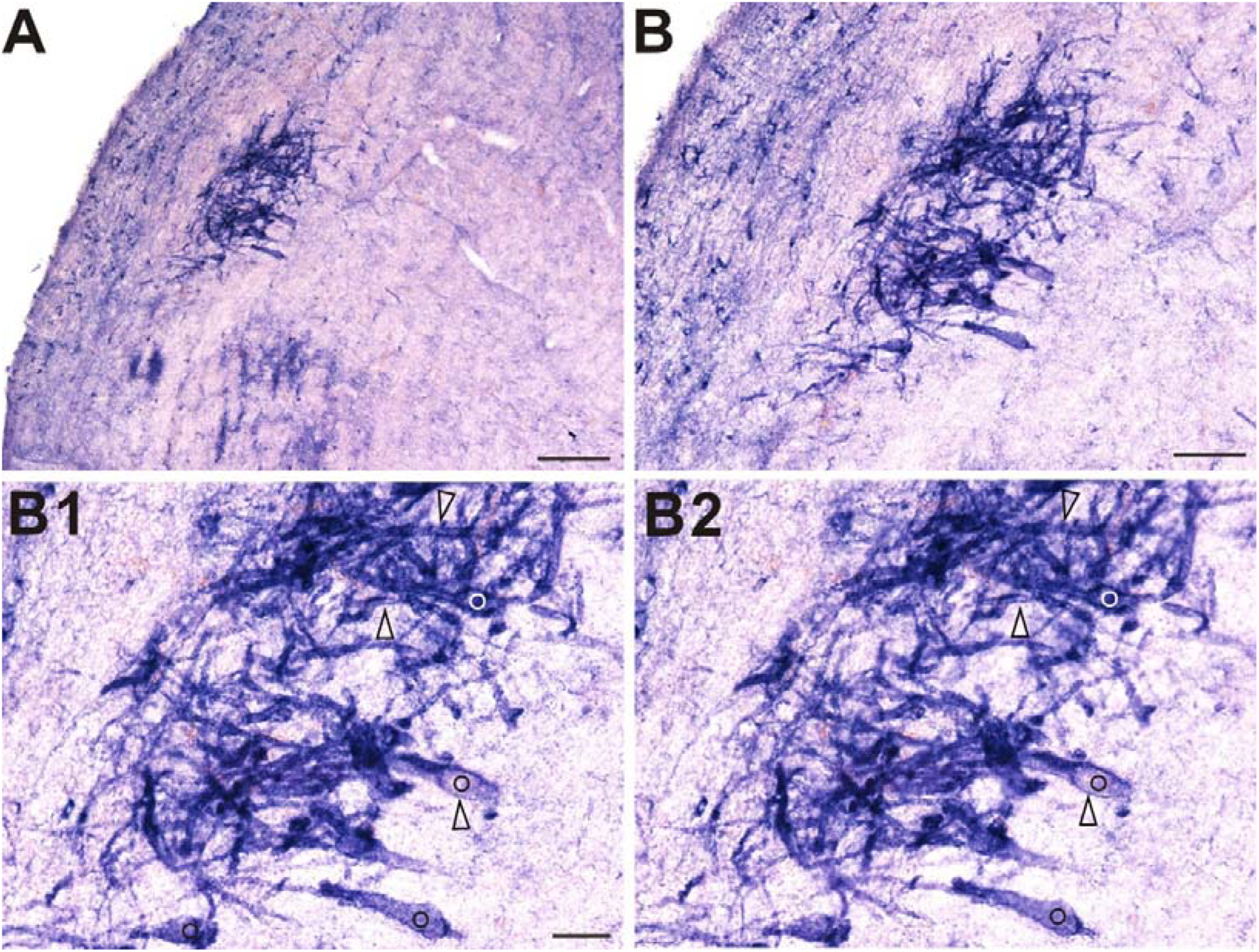
One example of N-d tubular glomerular structure in horizontal transverse section in the brainstem, with control enhance staining. A: The T-J body was located medial to the lemniscus spinalis, Bar = 100μm. B: Bar =50μm, B1 and B2: Higher power magnification of the same objects of two focusing images to show membrane-related N-d positivity viewed under the microscope. Three paired open arrowheads indicate three tubular parts in the two focusing images. Paired circles indicate perikaryons. Bar =20μm.

**Figure 4.**
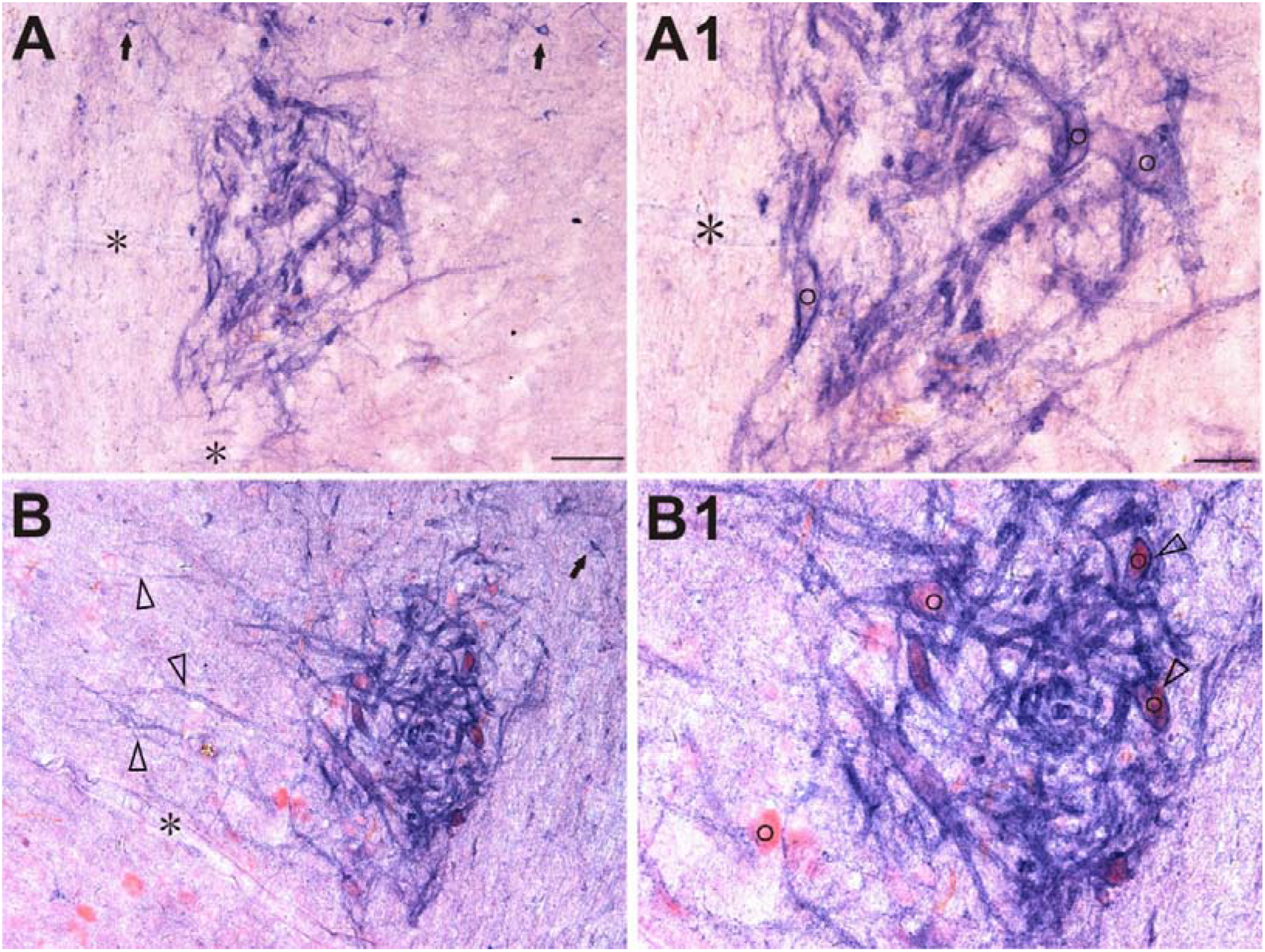
Peri-N-d-cytes and tubular glomerulus in horizontal sections with regular moderate N-d staining. A: N-d tubular glomerulus. Arrows indicate N-d intracellularly stained neurons. A1: Higher magnification of A. Asterisk indicates endothelium. Circles indicate perikaryon positions. B: Tubular glomerulus counterstained with neutral red. Open arrowhead alone indicates N-d positive and membrane-related tubule-like formation. B1: Higher magnification of B showed tubular glomerulus counterstained with neutral red. Circles indicate perikaryons with unstained N-d reactive intracellular formazans. Asterisks indicate endothelium. Bar for A and B = 50 μm. Bar for A1 and B1 =20μm.

**Figure 5.**
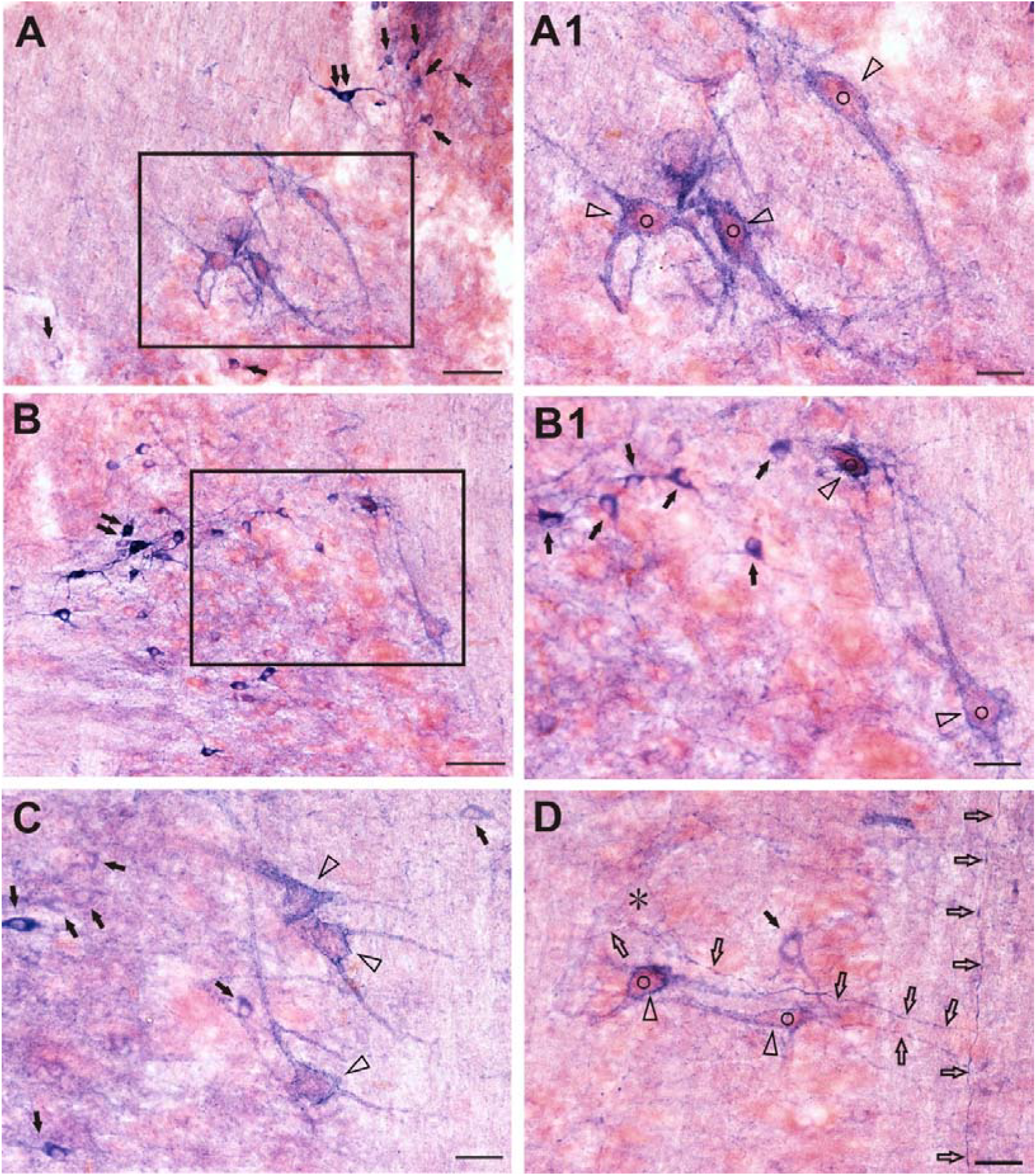
Large-sized peri-N-d-cytes with N-d positive proximal tubular processes revealed in the rostral to T-J body counterstained with neutral red in the horizontal sections. A: At least three Peri-N-d-cytes were showed rostral about 400-500μm to T-J body. Double arrows indicate strong reactive N-d neuron. Single arrows indicate various intensity N-d neurons. A1: magnification of A. Cirlce plus open arrawheads indicate Peri-N-d-cytes, which were multipolar neurons. B: Another example of Peri-N-d-cytes. At least three strong reactive neurons (double arrows) were showed high intensity formazans that covered nucleus. B1: magnification of B. Arrows indicate several small N-d neurons with intracellular N-d positivity. C: Three peri-N-d-cytes were showed, while several small neurons indicate by arrows. D: Two peri-N-d-cytes (circle + opened arrowhead) contracted nearby light staining neuron (arrow) showed perikaryon N-d positivity. Open arrows indicate fine N-d fibers. Asterisk indicates light stained endothelium. Bar in A and B = 50μm. Bar in A1, B1 C and D =20μm

**Figure 6.**
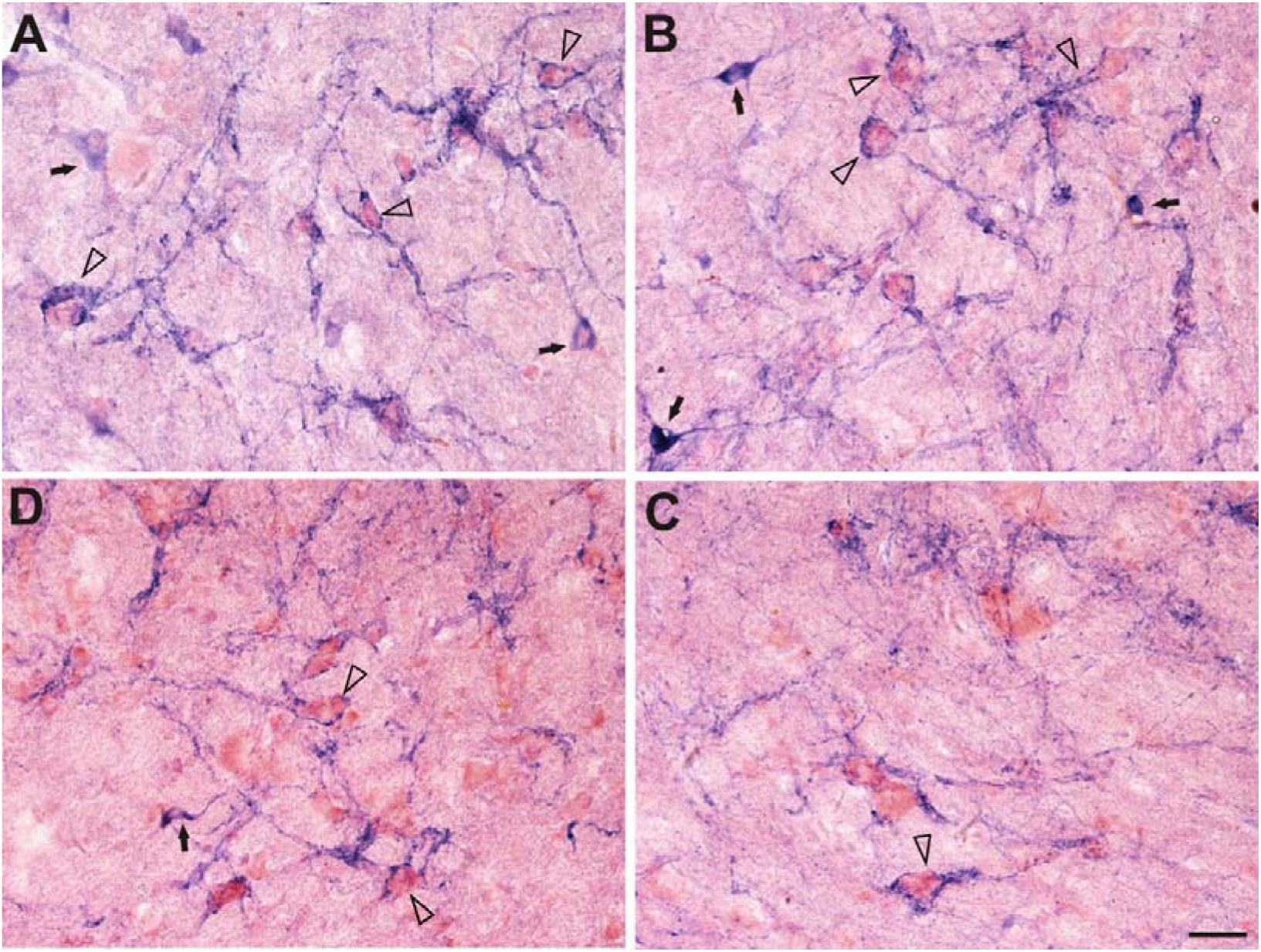
Another peri-N-d-cytes: patch N-d positive surrounding neurons in the rostral brain stem. Arrow indicates N-d neurons. Open arrowhead indicates peri-N-d-cyte. Some peri-N-d-cyte in A and B with thick proximal dendrites showed patched N-d positivity. Some N-d positivity incompletely surrounded the perikaryons. Bar =20μm.

## Discussion

The major finding of the present investigation is a new structure and N-d formazans surrounding unstained neurons by N-d histochemistry staining in the pigeon brainstem. Temporally, we termed it as N-d tubule glomerular structure. In order to easy to name it, we would like name it as T-J body as mentioned above. N-d formazans surrounding unstained neurons was termed as peri-N-d-cyte. In order to identification of the new finding, the phenotype classification of N-d neurons was demonstrated in several aspects. Various morphological formations were visualized by N-d staining. For the neuronal somas (perikaryon) and its fibers, several types of N-d positivity are classified: Golgi-like staining, strong staining intensity of soma with fibers, strong staining intensity of soma without clear distal positive fibers, weak staining intensity of soma with fibers or without distal fibers, projecting fibers (ascending, descending, cross or decussation), N-d positive puncta and varicosity. Beside capillary and or micro-blood vessels, the N-d tubular glomerular components in the T-J body was postulated as a unique formation of N-d positivity.

The gross cytoarchitecture of the pigeon’s medulla oblongata, stained with cresyl violet, was consistent to that observed in our recent study[8]. The basic morphological characteristics of N-d-positive neurons and fibers related to pigeon are demonstrated in several investigations[15, 17–19]. Atoji clearly investigates the distribution of N-d-positive neurons in the pigeon central nervous system[15]. We noted that the tubular components are not mentioned in their N-d mapping study[15]. In the other investigation of N-d positivity in the avian, we still found that no similar structure is demonstrated in the similar N-d mapping study in the chicken brain [20]. In adult budgerigars (Melopsittacus undulatus), the investigation of sexual dimorphism of vocal control nuclei is also not reported the similar ours finding[21]. Nevertheless, there was still existence of N-d neuronal cells among the tubular components, although the major part of T-J body was non-soma structures. The T-J body was also still revealed in the aged pigeons (data not showed here). The major morphological criteria of the T-J body in the aged animals was consistently with those of the young adults. However, the function of T-J body was not clear.

It is a specialized structure that could be functioned as a nucleus or a functional module or specific elements of neuronal parts. The enzyme N-d may not be only function for generation of neurotransmitter of nitic oxide. The bioactivity of N-d in the T-J body may indicate constructive structure for non-signaling function, because the N-d positivity formed tubular elements. And all the tubular components de-attached to perikaryons, so that N-d in T-J body may hardly function for neurotransmitter release from presynaptic terminals. Or how it functions release nitric oxide in such wiring device? T-J body bilaterally takes a large anatomical space in the pigeon brainstem. The large massive structure with such the N-d homogeneity unit is a potential device for specialized physiological function.

In order to further clarification of the T-J body, the related literatures were reviewed to demonstrate the relevant nueroanatomy of the medulla oblongata in pigeon or avian brain stem. The following investigations of the relevant anatomical location about the T-J body are discussed for a topographic analysis. The T-J body was rostral-caudal orientated and a longitudinal structure. The T-J body was located medial to the lemniscus spinalis, and ventral to several nuclei, such as nucleus angularis, nucleus et tractus tegmini descendens, pedunculo-pontius pars compacta, nucleus magnocellularis, nucleus parahrachialis and pars ventralis etc [20–26]. We cited following publications to help to localize the structure. Some nuclei such as superior olivary nucleus[27–29] and dorsal nucleus of the lateral lemniscus[28]could co-localize or overlap with T-J body. Or, a relative neighboring position could be plexus of Horsley or facial motor nucleus[27, 30–33]. We also noted that no similar T-J body was found in the chicken with Golgi staining[29]. According to our previous studies, a similar structure of the T-J body hasn’t been observed in rat, mouse, dog and monkey[6–8].

To our best knowledge, the moderate stained N-d tubular glomerulus structure is a specialized arrangement and formation. The characteristic of the N-d positive specialization was composed of tubular components bilaterally located in the medial to the lemniscus spinalis in the brainstem. The typical physical phenotype of the structure may implicate sounding analysis if it was related to olivary nuclei. It also hardly makes a postulation if our finding was assumed with plexus of Horsley for the time being. For testing the membrane location, enhanced N-d reaction did not show diffused formazans in the perikaryon’s area and inside tubular components (Figure 3). We found that the some of the peri-N-d-cytes were a large size perikaryons with large-diameter processes, which are considered for the faster velocity of impules[34, 35]. We thought that the existence of the similar peri-N-d-cytes could be reviewed in the nucleus magnocellularis cochlearis [15]. Where the N-d protein synthesis for the original derivation? How does N-d re-locate to the membrane from original produce site? We also do not know whether the peri-N-d-cyte worked as a nitric oxide provider or receiver. Working as a capacitor was another hypothesis for the N-d membrane-related location. Tubular units wired in the whole T-J body could be able to generate a special action by charging and or store electrical energy by synchronization of the signaling inputs. This very preliminary study needed more experiments to verify its identification.

## Conclusion

It could be a different intracellular and extracellular distribution for the N-d positive specialized structures featured with tubular components twisted and intermingled formation in the young adult pigeons. The peri-N-d-cyte and N-d membrane-related locations suggested that it may be new form of neuronal signaling transmission or a new morphology and functional constitution. The relevant of synaptic plasticity may also implicate some neuropathology.

## Supporting information

Supplement figure 1. Regular N-d positive staining of various somas and fibers revealed with diffused pattern in the brainstem. All of the phenotypes

## Acknowledgments

This work was supported by grants from National Natural Science Foundation of China (81471286), Undergraduate Training Programs for Innovation and Entrepreneurship of Liaoning (201410160007).

## Author Contributions

YJ and WH conceived and performed the experiments as well as analyzed data. WH, YL, XiaoxW, XinhW, TZ, HS, XianhW, LB, WZ, ZZ, JW, assisted YJ and HT in the experimentation. HT and YJ provided important experimental guidance. HT, YJ and ZZ discussed the results. HT wrote the manuscript. HT supervised the project and coordinated the study.

---

**Supplement figure 1.**
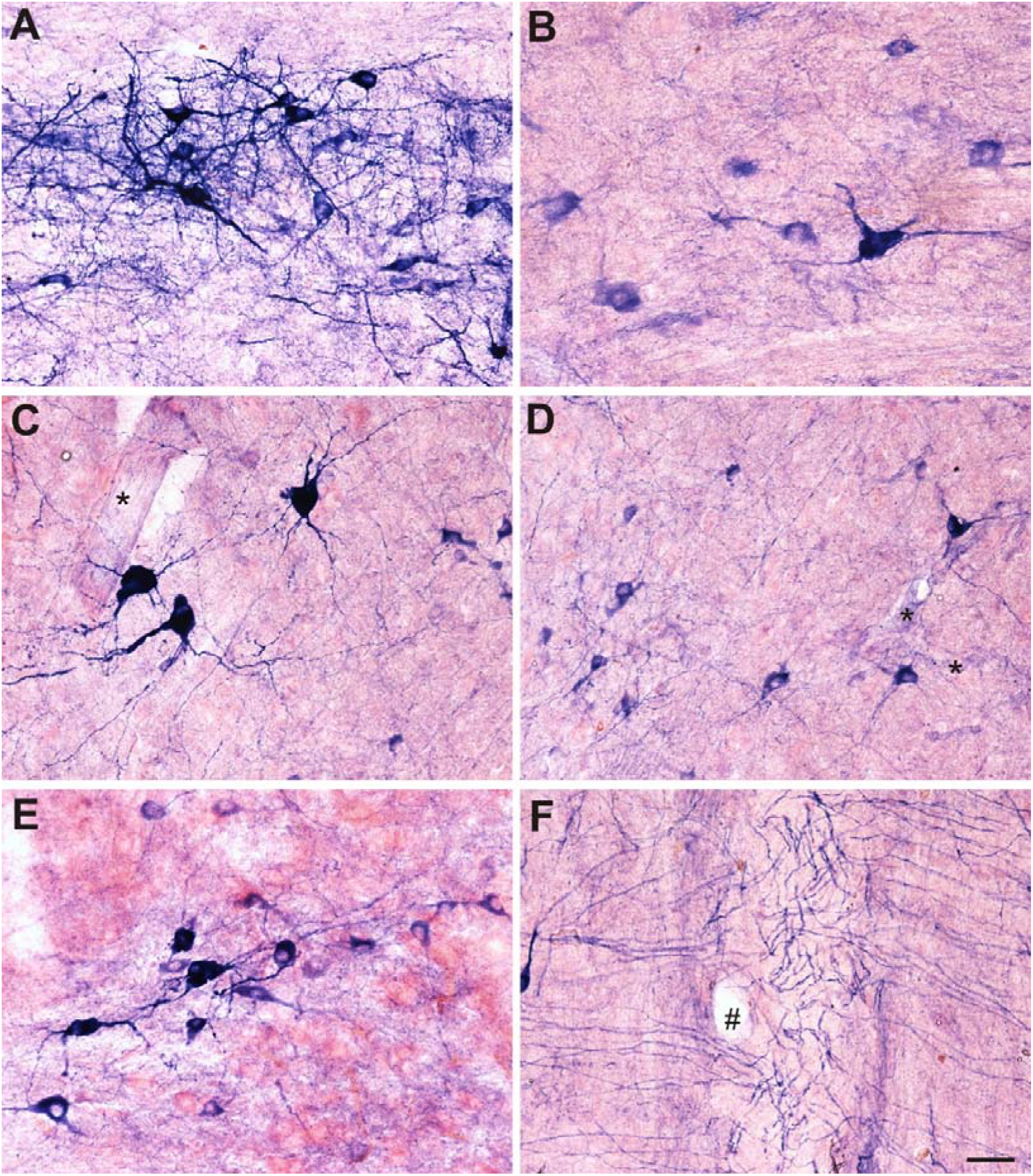
Regular N-d positive staining of various somas and fibers revealed with diffused pattern in the brainstem. All of the phenotypes of the positivity were observed in the horizontal sections. Formazans of N-d reactivity filled in most of the positive neuron in the whole intracellular area. Stronger reactivity masked nuclei of the perikaryons. The typical Golgi-like staining. Cross N-d fibers indicated varicosity. C-F: Sections were counterstained with the neutral red. All the images of the sections were procedure with the same incubation of the N-d staining. Asterisk indicates N-d positive endothelium of blood vessel. # indicates blood vessel. Bar: 20μm.

