## Supplementary figures and images for "Specialized NADPH diaphorase membrane-related localizations in the brainstem of the pigeons (Columba livia)"

### Supplement figure 1. Regular N-d positive staining of various somas and fibers revealed with diffused pattern in the brainstem. All of the phenotypes

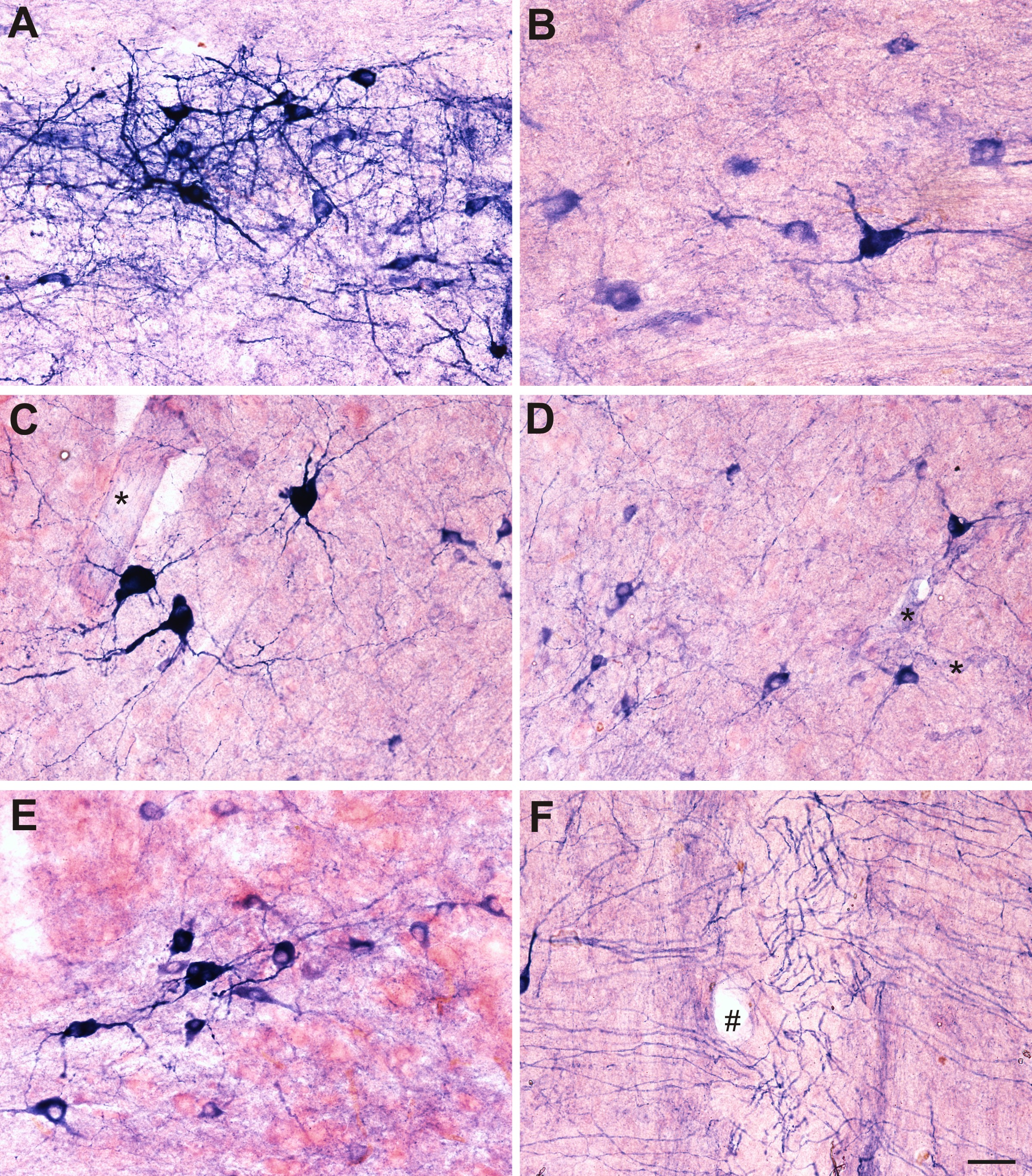
